# Effects of physiological deficits in pineal melatonin on Triple Negative Breast Cancer

**DOI:** 10.1101/2020.12.16.423028

**Authors:** Carley G. Dearing, Primal D. Silva, Danielle N. Turner, Betty A. Fish, Ethan O. Contreras, Zuri A. Schwarting, Johnny Sena, Faye D. Schilkey, Elaine L. Bearer, Stephen D. Wilkinson, Snezna Rogelj, Stewart Thompson

## Abstract

**Background:** Triple negative breast cancer (TNBC) is aggressive and treatment resistant. Evidence suggests that deficits in melatonin signaling increase TNBC risk: conditions that suppress melatonin increased incidence, low melatonin receptor expression correlates with worse prognosis, and high-dose melatonin can inhibit TNBC. Together this suggests that normalizing pineal melatonin could reduce TNBC incidence and/or mortality. The goal of this study was to determine whether small physiological deficits in melatonin alone, can increase risk for TNBC, and how ‘normal’ melatonin would be protective.

**Methods:** The effect of melatonin treatment on 4t1 cells *in vitro* was measured using the MTT cell viability assay, and gene expression of breast cancer and melatonin signaling markers. The effect of pineal gland status on 4t1 cell allografts was tested in C3Sn mice (*Mus Musculus*) with either an intact pineal (control) or surgical removal of the pineal causing a ~50% deficit in plasma melatonin. Orthotopic tumors were assessed by histopathology and metastasis by strain specific qPCR against 4t1 cell and host gDNA.

**Results:** Melatonin treatment induced significant changes in gene expression, with a significant reduction in derived PAM50 Risk of Recurrence score (ROR in Not treated = 65.5 ± 10.6 Mean SEM; Treated with 25pg/ml of melatonin 20.8 ± 8.3; P = 0.008), suggesting melatonin treatment would improve prognosis. A ~50% reduction in plasma melatonin increased orthotopic tumors, but this was non-significant, and had no effect on metastasis from tail vein allograft.

**Conclusions:** Physiological deficits in melatonin do alter the oncogenic status of 4t1 tumor cells but this has only a limited effect on growth and metastasis *in vivo*. Lack of significance in orthotopic tumor formation may be due to small sample size, and it is possible that any protective effect of melatonin occurs earlier in tumor development than we have tested.

## Introduction

Breast cancer is a leading cause of death among women in industrialized countries, with ~250,000 new cases and ~40,000 deaths each year in the United States [1]. The etiology of breast cancer is multifactorial and can result in a spectrum of presentations that have divergent prognoses. Notably, hormone receptor deficient breast cancers are typically aggressive and treatment resistant, with triple negative breast cancer (TNBC) accounting for 12-17% of cases, but having a higher proportion of total mortality [2].

Because TNBC is often treatment resistant and life-threatening, factors that affect incidence and/or severity can therefore be effective in limiting cancer morbidity and mortality [3]. One surprising risk factor for breast cancer is exposure to artificial light-at-night (ALAN). ALAN correlates with increased breast cancer incidence, and night shift work that would expose individuals to ALAN correlates with a marked increase in estrogen receptor negative breast cancer [4–7]. Consistent with this correlation, a reduced response to light in blind women has a protective effect against breast cancer [8,9].

Light acts on multiple aspects of physiology and behavior and the mediating mechanisms of these effects may be complex [10]. Current evidence does suggest that the suppression of pineal melatonin by light is a major contributor to these effects [11]. Pineal melatonin is suppressed by light at night in a dose dependent manner [12,13]. Additionally, melatonin receptors are expressed in breast tissue and MT1 melatonin receptor expression correlates with positive outcomes in breast cancer [14–16]. Finally, high dose exogenous melatonin inhibits mammary cell division *in vitro*, and mammary tumor growth *in vivo* [17–20].

Although most research points to an anti-estrogenic protective effect of melatonin, in patients with TNBC, melatonin receptor expression does correlate with survival, and polymorphisms of melatonin signaling correlate with incidence [16,21]. Further, an effect of melatonin on TNBC cells is also supported experimentally. *In vitro*, melatonin has an oncostatic effect on TNBC cell invasiveness and proliferation [22,23]. *In vivo*, extremely high doses of exogenous melatonin inhibit TNBC xenograft growth, and have oncostatic effects on tumor microenvironment, especially immune and angiogenic markers [24,25].

Based on this evidence, we might hypothesize that maintaining optimal plasma melatonin levels would reduce the burden of TNBC [11,16]. However, these studies do not show whether physiological deficits in endogenous melatonin levels, such as would occur with exposure to ALAN, are sufficient to increase risk for hormone receptor deficient breast cancer. Nor do they adequately demonstrate which stages in neoplastic development are affected by melatonin: initiation, promotion, progression and/or metastasis [26].

The goal of this study was to determine whether physiologically realistic deficits in endogenous melatonin impact the behavior of a hormone receptor deficient breast cancer, using the mouse 4t1 TNBC cell line [27–29]. We first assessed effects of melatonin *in vitro* on cell viability and then on expression of genes that give a prognostic prediction and assess potential mechanisms of melatonin action [30,31]. We then tested the effects of pineal melatonin on 4t1 cells *in vivo*. Pineal melatonin competent C3Sn mice were sham operated (Intact-control), or had the pineal gland tip surgically removed (PinealX) to generate a model of ~50% melatonin deficiency [12,32–34]. First, an orthotopic 4t1 cell allograft was used to assess the effect of pineal melatonin on viability and formation of solid tumors [35]. Second, a tail-vein 4t1 cell allograft was used to assess the effect of pineal melatonin on metastasis [36].

## Materials and methods

### Cells

4t1 cells are a mouse mammary tumor cell line with features of TNBC: low *Esr1*, absent *Pgr*, and normal or low *Errb2/Her2* [28,29]. Cells were sourced commercially (ATCC, Manassas, VA), and grown in Roswell Park Memorial Institute medium (RPMI; ATCC, Manassas, VA), with 10% fetal bovine serum (FBS; MidSci, Valley Park, MO) and 1% penicillin/streptomycin (Pen/Strep; HyClone, Logan, UT). There was a maximum of five passages before experimental use.

### Animals

Animal care and use was conducted in accordance with U.S. Public Health Service Policy on Humane Care and Use of Laboratory Animals and approved by the New Mexico Tech Institutional Animal Care and Use Committee. C3Sn.BLiA-*Pde6b*^+^/DnJ (C3Sn) mice were selected for use in this study because they produce pineal melatonin, are highly susceptible to breast cancer, and do not suffer from retinal degeneration like the background C3H/He mouse [32,33,37,38]. C3Sn were commercially sourced (Jackson Laboratories, Bar Harbor, ME) and then bred on site. Food and water were provided *ad libitum* throughout the study.

### *Experiment 1*. Effect of melatonin on 4t1 cells *in vitro*

To test whether melatonin treatment affects 4t1 cells *in vitro* we designed a custom gene expression assay, then tested how a physiologically relevant concentration of melatonin affected gene expression.

Reliable measurements of *in vivo* plasma melatonin in C3H background mice show a circadian rhythm with low levels during the day of ≤ 5 pg/mL and a 6 to 8 hour elevation at night of ~25 pg/mL [33]. Our test concentration of no treatment and ~25 pg/mL was intended to represent the maximal physiological range of plasma melatonin concentration.

Our goal with gene expression assessment was to reliably identify effects of melatonin on tumor status and to probe mechanisms of melatonin effect. Gene expression is an effective tool in prognosis prediction for human breast cancer. To more directly relate our data to modern practice in human cancer assessment, we included mouse homologs of the panel of targets used on the Prosigna^®^ PAM-50 test [30,31]. Critically, all fifty targets have a homolog in the mouse. To that panel, we added known and plausible mediators of melatonin effects on oncogenic status and transcripts associated with tumor initiation, development, and metastasis to specific tissues (**Supplementary materials S1**).

We first tested whether melatonin treatment would affect the number of viable cells available for mRNA extraction. Cells were grown as described above, treated with melatonin, then assayed in a standard MTT assay: 3-(4,5-dimethylthiazol-2-yl)-2,5-diphenyltetrazolium bromide (ThermoFisher, Waltham, MA) [39]. MTT is a colorimetric assay designed to test the metabolic activity of a culture. To prepare these assays, melatonin was dissolved in Dimethyl sulfoxide (DMSO) at a concentration of 1 mg/ml. The cells were plated at 4,000 cell/well in a 96-well microtiter plate and treated at concentrations ranging from 200 pg/ml down to ~0.5 pg/ml. The cells were incubated for 48 h in 200 μl of RPMI media with 1% Pen/Strep and 10% FBS. Twenty percent v/v of MTT reagent in 1X PBS (5 mg/mL) was added to each well and incubated further for 2 h. Media was removed and replaced by 100 μl of DMSO. Absorbance at 595 nm was measured using a Thermomax Molecular Device plate reader. The experiments were performed in quadruplicate. A 0.1% DMSO was used as a vehicle control and 10 μM phenyl arsine oxide (PAO) was used as a positive killing control. Statistical analysis was limited to the dose response data for melatonin treated groups with Not Treated control assumed to be 0.0 pg/ml; assessment was by Welch’s ANOVA in Prism (GraphPad, San Diego, CA).

Then to determine how melatonin affects gene expression in 4t1 cells, we applied melatonin to cells *in vitro*, isolated RNA, and assessed expression of a custom panel of target genes.Cells were grown to confluence. Melatonin was added to the media in flasks at 25.0 pg/mL (N = 6 flasks) with no treatment controls (N = 6). After 4 hours, a sample of media was frozen for melatonin content quantification, then cells were dissociated using trypsin, stabilized in RNAprotect cell reagent (Qiagen, Germantown, MD) and frozen for later mRNA isolation.

For control tissue, 16-week old virgin female C3Sn mice were euthanized by anesthetic overdose (200 mg/kg Ketamine, 20 mg/kg Xylazine) at the circadian phase of peak melatonin production (6 to 9 hours after lights off) under far red light (720 nm). Blood was drawn and serum isolated for melatonin quantification, protecting melatonin content by minimizing exposure of the sample to light, and by storing at −80 °C after serum separation. The thoracic and abdominal lobes of the right breast were pooled and snap frozen for RNA isolation.

RNA was isolated using an RNAeasy Protect Cell Mini Kit (Qiagen) for *in vitro* cell samples, and an RNAeasy Mini Kit (Qiagen) for C3Sn mouse normal mammary tissue. All samples were reverse transcribed using an RT2 First Strand kit (Qiagen). Our panel of 88 target genes was prepared on a 96-well plate format with housekeeping genes and controls (RT2 PCR array, Qiagen). Expression was quantified on an ABI7500 real-time PCR machine (Applied Biosystems, Beverly, MA) using RT2 SYBR Green ROX qPCR mastermix (Qiagen). One non-treated sample was excluded for failing quality controls.

Gene expression data for the PAM50 panel genes was used to generate a breast cancer Risk of Recurrence (ROR) prognosis score [30]. The *“rorS”* algorithm in genefu version 2.23.0 01/31/2020 was applied [40]. The derived score is on a scale of 0–100: a *low* score (<40) indicates a 10-year ROR less than 10%, an *intermediate* score (40 to 60) indicates a 10-year ROR of 10–20%, and a *high* score (>60) indicates a 10-year ROR of >20%. Comparison of ROR scores in not-treated controls and cells treated with 25pg/ml of melatonin was by unpaired parametric 2-tailed t-test in Prism (GraphPad).

Then, data was normalized by CT of Symplakin (Sympk), which had the most consistent expression across samples (Mean CT 25.5 SD 0.7), and has been identified as an optimal housekeeping gene for breast cancer gene expression [41]. Percent change was calculated relative to mean expression in ‘not treated’ controls using a power of 2. Statistical significance of change in gene expression was determined by a two-sample equal variance two-way t-test.

### *Experiment 2*. Orthotopic allograft of 4t1 cells in C3Sn mice

A preliminary study of orthotopic allograft was conducted to demonstrate that BALB/c 4t1 cells could form viable tumor cell colonies in our melatonin competent C3Sn strain host [12]. This small sample size experiment also provided a preliminary assessment of whether pineal melatonin affected the formation of primary tumors and metastasis from those primary tumors.

#### Pineal surgery

Between 7-10 weeks of age, C3Sn mice were surgically pinealectomized (PinealX) or were sham operated leaving the pineal gland intact (Intact-control) [34]. To prevent selection and treatment bias, we evenly and randomly assigned animals in a litter to pinealX or sham surgery, then housed those animals together irrespective of their pineal status and masked experimenters to pineal status. Analgesia and anesthesia protocols included 100 mg/kg Ketamine; 10 mg/kg Xylazine; 2.5 mg/kg Acepromazine; Buprenorphine HCL at 0.1 mg/kg; and post-surgical access to an oral tablet containing 2mg of the NSAID, carprofen (Bio-Serv, Flemington, NJ). After surgical site preparation, mice were placed on a heat mat and mounted into a stereotaxic frame (Kent Scientific, Torrington, CT). A 1.5 cm midline vertical incision and blunt dissection was used to expose the scalp. A section of skull, centered on the intersection of the sagittal and occipital fissures of the calvarium was removed using a hand drill and 2.3 mmø trephine drill bit (Fine Science Tools, Foster City, CA). This approach removes the tip of the pineal gland with the cap of skull that is removed but should leave part of the pineal intact. Removal of the pineal gland was confirmed using a SZ-745 dissecting microscope (McBain, Westlake Village, CA), and the incision closed with non-absorbable 5-0 sutures (Ethicon, San Angelo, TX).

#### Allograft

After 2 weeks of surgical recovery, mice had an allograft of 4t1 cells to the fat pad of the left 4th mammary gland using a previously described approach [42]. Cells and mice were prepared in parallel to minimize time between cell preparation and allograft (N for each group = 8, 4 male and 4 virgin female). 4t1 cells were dissociated with trypsin in RPMI media, then suspended in RPMI at a concentration of 4*10^7^ cells/ml. Within 60 minutes, the 4t1 cell preparation was injected orthotopically to the 4th left mammary fat pad of mice. Mice were lightly anesthetized with 50 mg/kg Ketamine; 5 mg/kg Xylazine; 1.25 mg/kg acepromazine. The surgical site was prepared and a small incision made approximately 1.5 mm above the 4^th^ nipple. 50 μl of 4T1 cells totaling 2*10^6^ cells was then injected into the fat pad using a gas-tight micro-syringe (Hamilton, Reno, NV). Because the incision was small and shallow, no suture was used to close it.

#### Tumor development and assessment

After recovery from allograft anesthesia, mice were housed in environment control cabinets under a daily cycle of 12-hours light (20 μWcm^-2^), and 12-hours dark. At 16 weeks post cell injection, the mice were euthanized by anesthetic overdose followed by cervical dislocation. Hair was removed from the abdomen of the mouse using depilatory cream (Nair, Ewing, NJ). Left and right mammary chains and the lungs of each animal were collected and fixed in 4% paraformaldehyde (PFA) (Sigma-Aldrich, St. Louis, MO) for 6 hours, then transferred to 1% phosphate buffered saline (PBS) and stored at 4 °C. Tissue was embedded in Tissue Freezing Medium (General Data, Cincinnati, OH) sectioned at 10-16 μm on a Shandon FE Cryostat (Thermo Fisher scientific, Waltham, MA). Slides were stained with Hematoxylin and Eosin (all reagents from VWR, Radnor, PA). Number of sections was recorded to allow size calculation. Images were recorded on a Leica ICC50 HD and processed with Microsoft Image Composite Editor.

Larger tumors were defined as a cluster of dense cells with a defined border demarcating it from surrounding normal adipose tissue. We assumed a solid tumor was a single mass unless there were three sequential sections with no tumor between two masses. After assessments, animal pineal status was unmasked and total tumor load compared between experimental PinealX and Intact-control groups using Welch’s *t*-test.

Single epithelial cells in the fat mass of the mammary pad might be invasive 4t1 cells. To assess differences in this potential marker of local invasiveness, we quantified the extent of triple negative tumor presence in the left breast away from the introduction site at the abdominal or 4^th^ mammary gland using a percentage-based severity grade. Invasiveness was defined as a small dense cluster of cells that displayed branching into surrounding tissue. The invasive burden was measured by counting invasive clusters of cells in a section (0-7, 7 being the most severe). Because this scoring approach is subjective, scorers were masked to animal ID and analysis by three separate scorers was averaged. After assessments, animal pineal status was unmasked and total tumor load compared between experimental PinealX and Intact-control groups using Welch’s *t*-test.

### *Experiment 3*. Effects of pineal melatonin on metastasis of 4t1 cells from tail-vein allograft

It has been demonstrated that tail-vein injection allograft of 4t1 cells establishes metastases [36]. We therefore adopted this model to determine whether melatonin affects the extent and sites of metastases of 4t1 TNBC cells. We focused on common metastatic sites for breast cancer: breast, liver, and lymph nodes. However, there was potential for low levels of metastatic burden in these tissues, so we developed a gDNA based quantification host and donor content in tissues using strain selective qPCR (**see supplementary material S2**). This is similar to the approach used in quantifying xenografts by others [43–45].

#### Pineal surgery

Mice were Intact-control (n=10, 6 female and 4 male) or PinealX (n=13, 6 female and 7 male) operated between 7 and 10 weeks old as described for experiment 3.

#### Tail-vein injection

After 2 weeks of surgical recovery, mice had an allograft of 4t1 cells to the tail-vein using a previously described approach [27,36]. Cells and mice were prepared in parallel to minimize time between cell preparation and allograft. 4t1 cells were dissociated with trypsin in RPMI media, then suspended in RPMI at a concentration of 5*10^7^ cells/ml. Within 60 minutes, 0.1 ml (500,000 cells) were injected into the tail vein of mice lightly anesthetized with 50 mg/kg Ketamine; 5 mg/kg Xylazine; 1.25 mg/kg acepromazine. After cell injection, mice were housed in a light tight environment control cabinet with a 12-hour light, 12-hour dark daily cycle.

#### Tissue collection

At 14 days post allograft, mice were euthanized by anesthetic overdose (200 mg/kg Ketamine, 20 mg/kg Xylazine) at the circadian phase of peak melatonin production (6 to 9 hours after lights off) under far red light (720 nm). Blood was drawn and serum isolated for melatonin quantification, protecting melatonin content by minimizing exposure of the sample to light, and by storing at −80 °C after serum separation. Tissue samples of common metastatic sites were then collected into dry tubes for snap-freezing. Each lobe of the lung was collected separately: right cranial, middle, caudal, accessory and left lung. Both inguinal lymph nodes were pooled. The thoracic and abdominal sections of the right breast were pooled. Liver collection was limited to a ventral biopsy.

#### Melatonin status confirmation

A direct melatonin immunoassay was used to measure melatonin concentration in serum at the time cells were harvested (MEL31-K01 direct melatonin serum/plasma EIA kit, Eagle Biosciences, Nashua, NH). Samples were tested according to manufacturer guidelines and absorbance at 450nm measured on an Infinite MPlex 96-well plate reader (Tecan, Baldwin Park, CA). A standard curve was plotted using a 4-parameter sigmoid function, and concentrations interpolated from the curve, in Prism (GraphPad).

#### DNA isolation

DNA was isolated using a DNeasy Blood and Tissue kit (Qiagen). After adding a lysis buffer, tissue was mechanically disrupted using a Tissueruptor II with disposable probes (Qiagen). Further lysis was promoted with Proteinase K step for 2 hours before finishing DNA isolation. The exception to this protocol was for bone, where the Proteinase K step was 40 hours, and then mechanical disruption was completed with a Tissueruptor II. Quality of DNA was tested by 260/280nm or 260/230nm ratios on a Nanodrop spectrophotometer (Thermo Fisher). Quantity of DNA was measured by the 260nm adjusted absorbance.

#### gDNA quantification

Isolated gDNA was amplified in triplicate with strain-selective BALB/c and C3H primers (IDT, Coralville, IA) and a qPCR and Go SYBR^®^ Hi-ROX Kit (MP Biomedicals, Irvine, CA) on an ABI7500 real-time PCR machine (Applied Biosystems). Threshold cycle (C_T_) values were converted to percent of total gDNA using a power of 2. After assessment, animal pineal status was unmasked and total tumor load compared between Intact-control and PinealX groups using a two-tailed equal variance *t*-test.

## Results

### *Experiment 1*. Effect of melatonin on 4t1 cells *in vitro*

There were viable 4t1 cells under all concentrations of melatonin treatment, so we were able to select a concentration of melatonin for testing gene expression that was a maximal physiological concentration (**Figure 1**). Melatonin had no identifiable effects on the gross morphology of 4t1 cells. In addition, there was no effect on proliferation/survival of 4t1 cells in an MTT assay (Welch’s ANOVA of melatonin treatment dose response curve P = 0.29; W 1.4, DFn 9.0, DFd 11.3). Comparison of no treatment control confirmed that there was no difference to DMSO vehicle control (Mann Whitney test P = 0.20), and that cells were effectively killed by PAO positive control.

**Figure 1.**
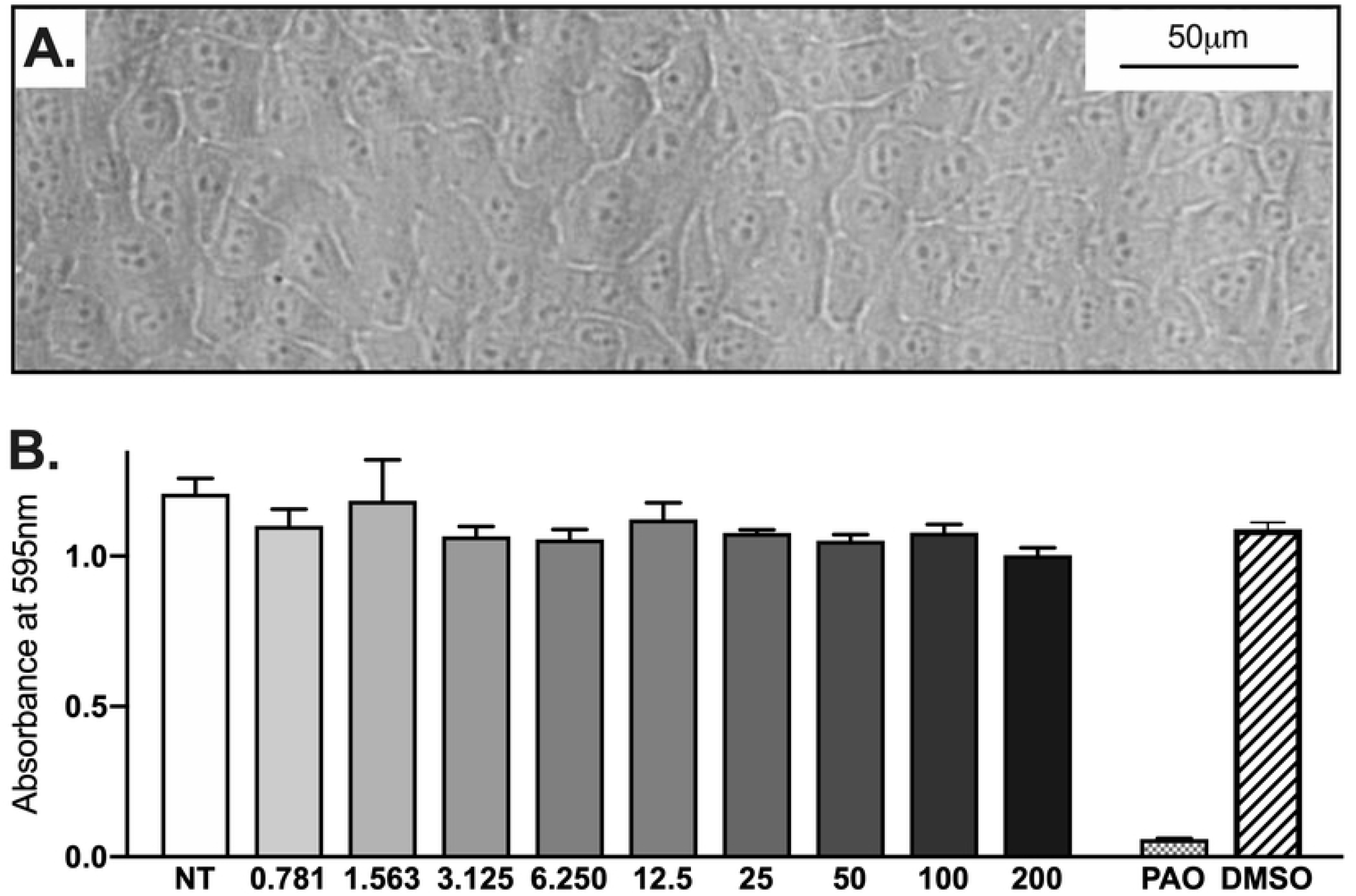
Effects of melatonin on 4t1 cell morphology and viability. **(A)** An example image of 4t1 cells with melatonin treatment at a concentration much higher than physiological levels. **(B)** In the MTT assay, viable cells breakdown of tetrazolium dye to formazan, which has absorbance at 595nm, so absorbance = metabolic activity = viable cells. Absorbance for 4t1 cells is shown with melatonin treatment at a range of concentrations. Melatonin concentration gradient is implied by shading of bars on the graph. PAO positive killing control and DMSO vehicle control are also shown. Data are expressed as mean ± SEM.

Gene expression confirmed the TNBC status of 4t1 cells, with significantly reduced expression of *Pgr* (<0.01% of normal breast expression, P< 0.00001), *Esr1* (1.2% of normal breast, P< 0.00001), and *Errb2/Her2* (10.4% of normal breast, P< 0.0001) **(Figure 2A, Table S1)**. These data also identified significantly lower expression of melatonin receptor *Mtnr1a* (3.0% of normal breast, P< 0.001), and the melatonin receptor regulated transcription factor *Rora* (0.05% of normal breast, P< 0.00001).

**Figure 2.**
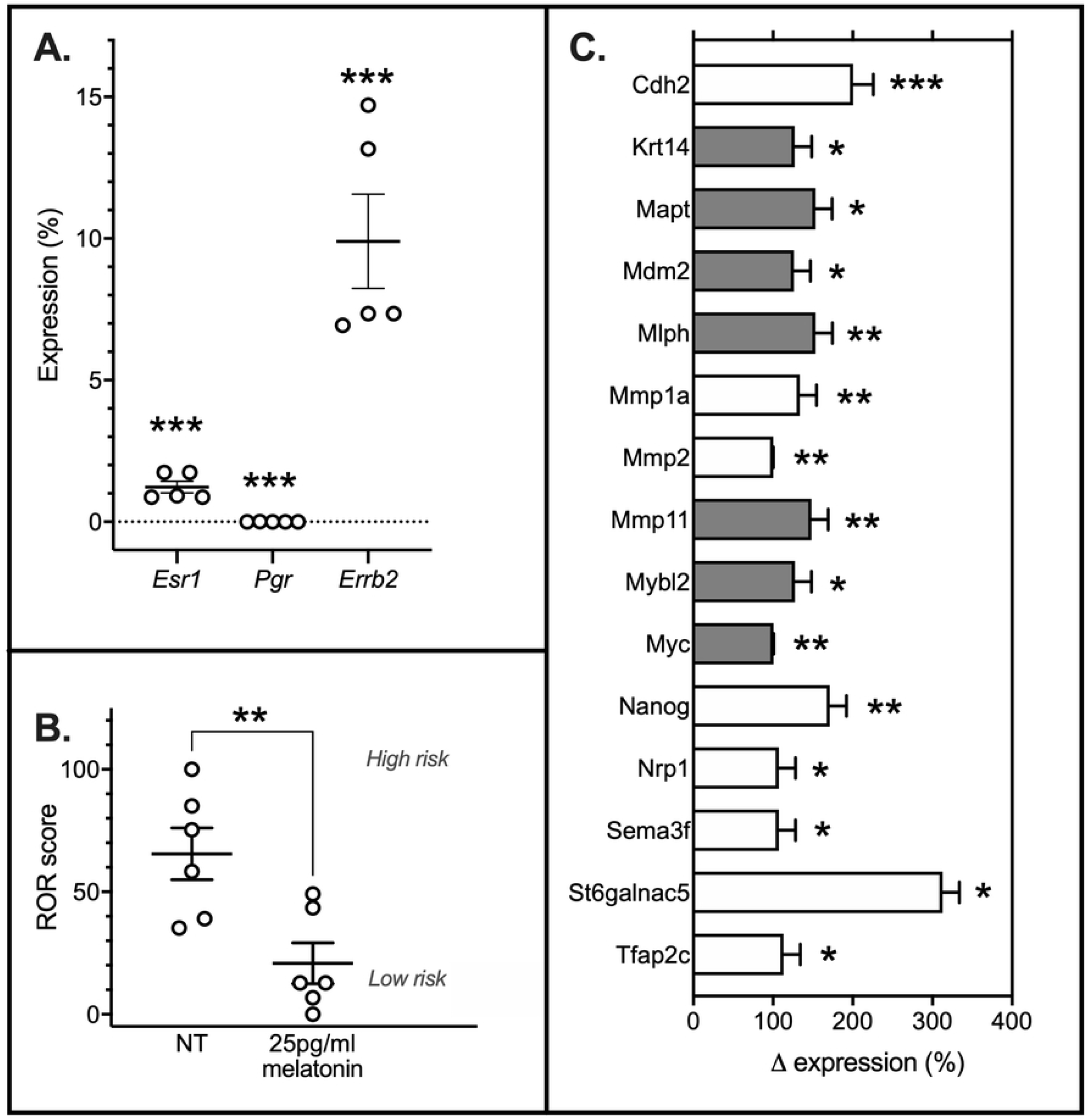
Gene expression of 4t1 cells *in vitro*. Expression of selected genes was determined by qPCR for 4t1 cells grown in cell culture media. **(A)** *Esr1, Pgr* and *Errb2/Her2* are the hormone receptors used to identify breast cancer as triple negative. Gene expression is shown as percentage of the expression in normal mammary tissue from C3Sn mice, after normalizing samples to a housekeeping control *Symplekin*, with bars showing Mean and SEM. **(B)** Expression levels of the PAM50 panel of genes allowed a Risk-Of-Recurrence score (ROR) to be calculated for experimental replicates of 4t1 cells grown in untreated cell culture media (NT) and in media treated with 25pg/ml of melatonin for 4-hours, which approximates peak plasma melatonin in mice. **(C)** Melatonin treatment significantly changed gene expression of multiple targets in our assay. Shading shows genes that are part of the PAM50 panel, non-shaded genes were our melatonin mechanism focused additions to the panel. Mean and SEM percent change was calculated against expression in cells with no melatonin pretreatment. Significance in all panels is indicated by * P<0.05, ** P<0.01, and *** P<0.001.

There was an effect of melatonin treatment within the physiological range (25pg/ml) on gene expression of 4t1 cells in culture **(Figure 2B)**. Assessment of the genes that constitute the PAM50 human breast cancer Risk-Of-Recurrence (ROR) prognosis score calculator identified a significant decrease in oncostatic status with melatonin treatment (P = 0.008; F = 1.61). Non-treated cells had a *high* ROR score (Mean 65.5 ± 10.6 SEM), which in human patients identifies 10-year ROR as greater than 20%. Cells treated with 25pg/ml of melatonin had a *low* score (Mean 20.8 ± 8.3 SEM), which in human patients identifies 10-year ROR as less than 10%.

Among the PAM50 panel and our added targets, fifteen genes showed a significant increase in expression with melatonin pretreatment **(Figure 2C).** These included genes that: anchor cells into tissue (*Cdh2, Nrp1* and *Sema3f*), enable invasion and metastasis (*Krt14, Mlph, Mmp1a, Mmp2, Mmp11, St6galnac5*), and that affect the rate of differentiation, proliferation and/or apoptosis (*Mapt*, *Mdm2*, *Mybl2*, *Myc*, *Nanog*, *Tfap2c*). *Note*: when identifying changes caused by melatonin treatment, we did not correct for multiple measures. This approach is likely to include type 1 errors (false positives), but with Bonferroni correction, type 2 errors are likely (false negatives), and due to the large number of targets in our panel, with correction none of the changes were significant.

### *Experiment 2*. Effect of pineal melatonin on a 4t1 cell orthotopic allograft into C3Sn mice

Orthotopic allograft was intended to test whether 4t1 cells were viable in a C3Sn mouse, and to allow us to assess whether pineal melatonin had any effect on 4t1 tumor burden, invasiveness and metastasis from an orthotopic placement.

A tumor cell burden was present in the majority of mice, demonstrating that 4t1 tumors will establish themselves successfully in a C3Sn host mouse, regardless of sex or pineal/melatonin status **(Figure 3)**. There was a non-significant increase in the incidence of solid tumors in female PinealX mice (P = 0.38, F = 8.0). There were also single epithelial cells in the fat mass of the mammary pad that might be invasive 4t1 cells. If these are invasive 4t1 cells, there was no difference in the severity of local invasion between female PinealX and Intact-control (P = 0.68, F = 15.5).

**Figure 3.**
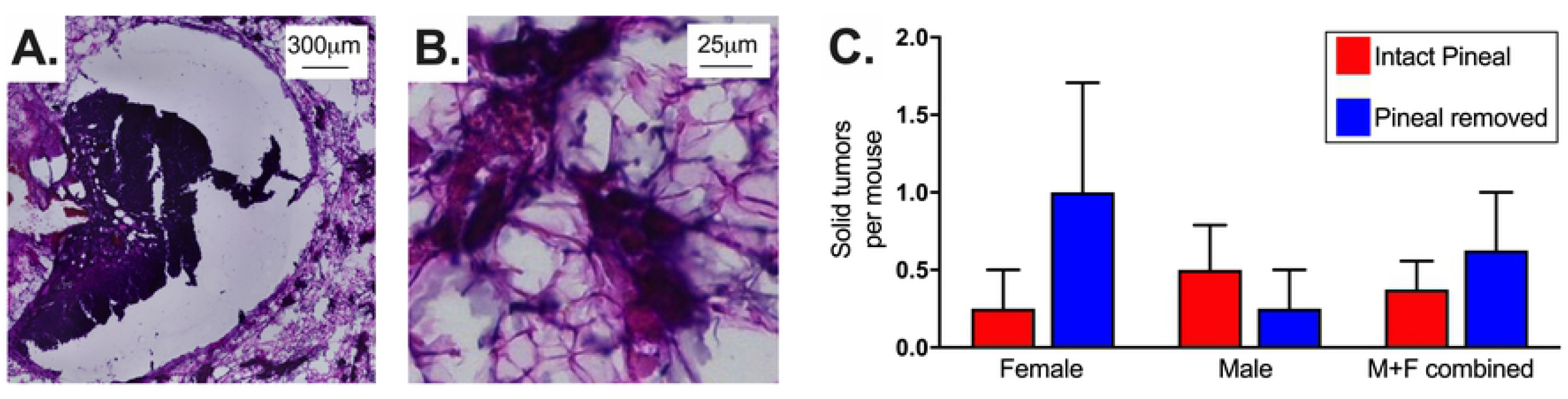
Effect of pineal melatonin on orthotopic allograft viability. **(A)** Example of a solid tumor at the allograft site in the lower left breast chain. **(B)** Example of single epithelial cells in mammary fat pad that might be invasive 4t1 cells. **(C)** The number of solid tumors in the lower left mammary chain of Intact-Pineal control and PinealX mice is expressed as Mean ± SEM. Tumor numbers are shown for female mice, male mice, and combined groups of male and female mice.

### *Experiment 3*. Effect of melatonin on metastasis of 4t1 cells from tail-vein allograft

There was no effect of pineal status on metastases to lung, breast, lymph or liver **(Figure 4)**. Pineal surgery significantly reduced serum melatonin (Mean ± SD serum melatonin pg/ml: Intact-Control 32.2pg/ml ± 4.7; PinealX 16.2 ± 2.1; P < 0.0001; F = 5.11). There was no sex difference in mid-dark-phase serum melatonin (Intact-Control Male 31.0 ± 5.8; Intact-Control Female 33.5 ± 4.0; P = 0.40).

**Figure 4.**
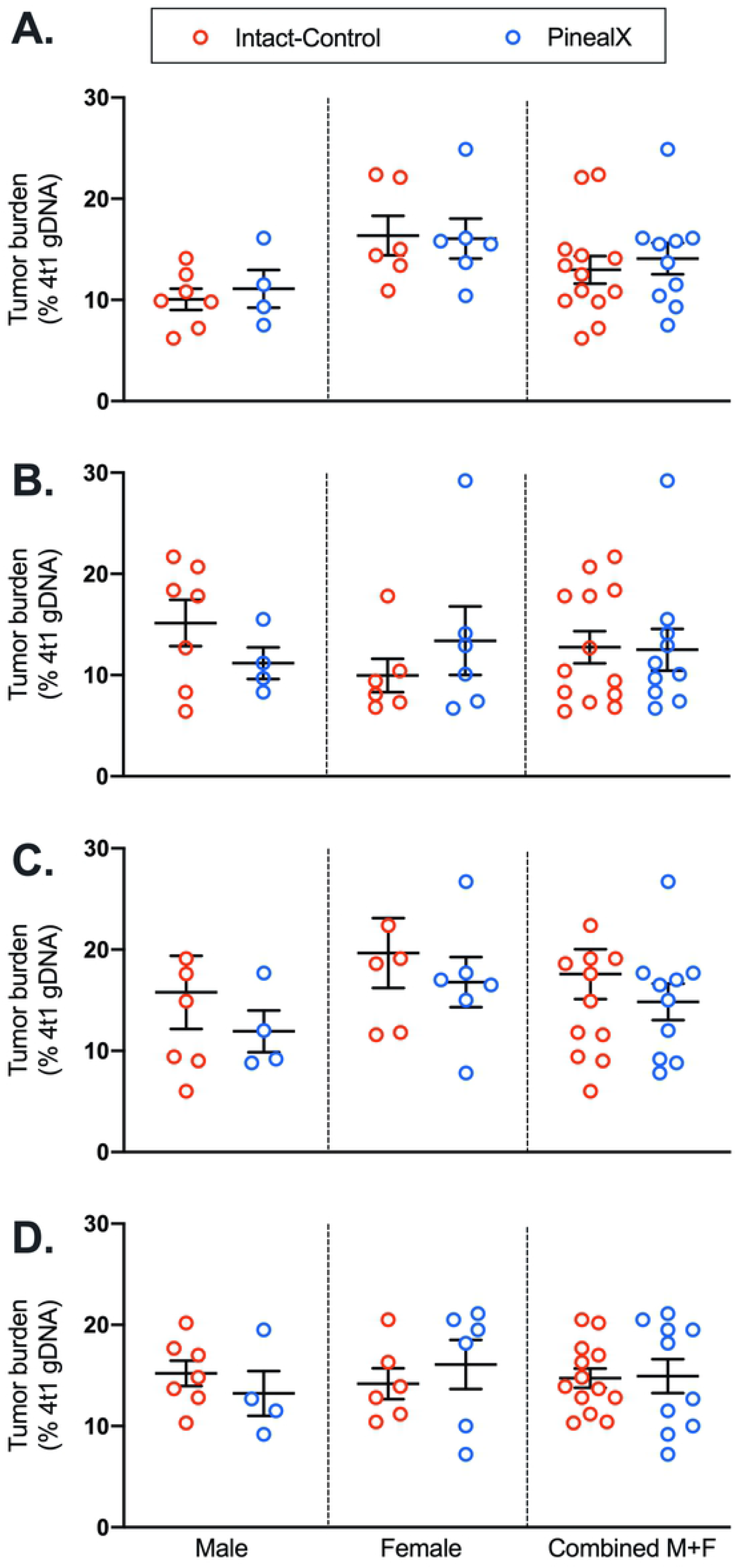
Effect of pineal status on metastasis. 4t1 cell burden is shown as a percentage of the total tissue calculated by strain specific qPCR: 4t1 cells have a BALB/c origin so the proportion of BALB/c and C3Sn host gDNA provided a measure of 4t1 cell content. Mean and SEM of data are shown for Intact-Control and PinealX mice for: **(A)** breast, **(B)** lung, **(C)** lymph, and **(D)** liver. Comparison is Intact-Control versus PinealX for male mice, female mice, and then combined male and female data.

However, there was a significant difference in metastases of 4t1 cells to the breast of male and female mice (Mean and SD females = 16.2 ± 4.6, males = 10.4 ± 3.0, P = 0.002; two-tailed, equal variance t-test, significance threshold Bonferroni corrected for multiple measures from 0.05 to 0.0127) **(Figure 5)**. There was also no effect of sex on metastases to lung, lymph or liver.

**Figure 5.**
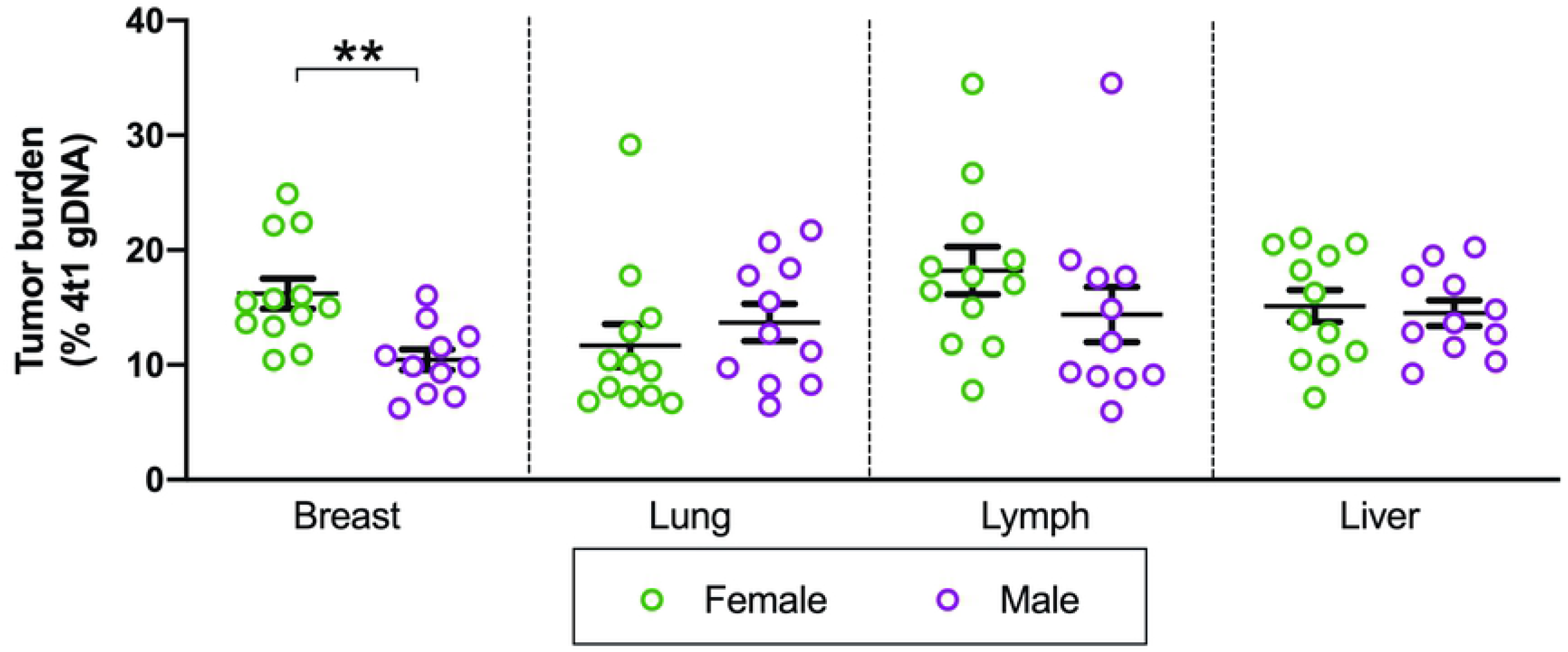
Effect of sex on metastasis. 4t1 cell burden is shown as a percentage of the total tissue calculated by strain specific qPCR. Mean and SD of data separated by sex are shown for: **(A)** breast, **(B)** lung, **(C)** lymph, and **(D)** liver.

## Discussion

There is compelling evidence that reduced melatonin signaling correlates with increased incidence and poor prognosis in TNBC patients [5,9,22,24,25,29]. However, studies of melatonin effects on TNBC have typically been limited to very high, non-physiological doses of melatonin, and/or used strains of mice that do not make endogenous melatonin so are chronic deficits in melatonin with unspecified developmental consequences of that deficit. The goal of this study was to determine whether the relatively small reductions in pineal melatonin that might occur with use of artificial light-at-night (ALAN), have any effect on a hormone receptor deficient breast cancer.

Assessment of the effect of melatonin on 4t1 cells *in vitro* was intended to identify any effect on tumor cell status (ROR score) and suggest mechanisms mediating effects of melatonin. The lack of melatonin effect on survival of 4t1 cells *in vitro*, was consistent with one study of MDA-MB-231 and HCC-70 TNBC cells, but different from a later study of MDA-MB-231 cells [22,23]. However, there was a clear effect of melatonin on gene expression. Gene expression also confirmed that 4t1 cells are valid as a model of TNBC: *Pgr* was undetectable, and *Esr1* and *Errb2/Her2* were considerably reduced. This is consistent with the finding that 4t1 cells lack Esr1 protein and an estradiol growth response [27]. However, this is different from the observed expression of *Esr1* and *Errb2* in another recent study, which likely reflects our use of C3Sn mouse to provide normal breast gene expression data [29].

Notably, the significant reduction in PAM50 ROR prognosis score with melatonin treatment suggested that melatonin has the potential to dramatically reduce the aggressiveness of 4t1 TNBC [30,31]. The caveat with the ROR observation is that the PAM50 panel and ROR prognosis score are developed and clinically validated for assessment of human breast cancer. However, we felt this panel would make both typing and prognosis assessment more translationally relevant in this and future studies. For example, the PAM50 panel was developed for a variety of breast cancer types (Luminal A, Luminal B, HER2-enriched, basal-like and normal-like), and tools to derive the clinically validated prognosis score as a 10-year risk of recurrence (ROR) are freely available [40].

Of the melatonin induced changes in specific genes, patterns of interest included an increase in genes associated with tissue remodeling (*Mmp1a, Mmp2, Mmp11*), but also an increase in cell-cell anchoring genes (*Cdh2, Sema3f, Nrp1*). The sum of these changes may be negative, positive or neutral, which emphasizes the value of a clinically validated prognosis predictor such as the PAM50 ROR.

Given the effect of gene expression, the limited effect of deficits in melatonin *in vivo*, was unexpected. We had successfully developed a surgical model of partial pineal gland removal that reduces plasma melatonin by ~50%. In our preliminary study to test allograft viability, we saw a non-significant increase in burden of solid tumors in female mice from 4t1 cells introduced orthotopically. However, in a larger cohort with 4t1 cells introduced by tail-vein injection, we saw no effect of pineal status on metastasis. Interestingly, the burden of 4t1 cells from tail-vein allograft was significantly higher in the breast of female mice. This measurement was proportional to total mammary tissue collected so could result from a sex-hormone effect on 4t1 cells or a functional difference in the mammary tissue of virgin female mice [46].

While we saw no effect of pineal melatonin on metastasis, our approach of quantifying metastases by gDNA was successful. Metastases of 4t1 cells from tail vein injection form disseminated colonies, which would make quantification difficult [36]. Others have developed methods for quantification of submicroscopic xenograft metastases using DNA quantification [45]. The genetic differences between BALB/c 4t1 cells and our C3Sn host allowed us to design strain specific qPCR primers, which allowed us to quantify dispersed metastases from a transplant to a member of the same species [47]. To our knowledge, this approach has not been applied to an allograft.

Our gene expression data suggests that melatonin does have a meaningful effect on TNBC. The lack of melatonin effect in an MTT assay and on metastasis from tail-vein injection, suggests the effect is not on proliferation or invasion of tissues from the vasculature. In that context, the trend for increased tumor burden from orthotopic allograft in females with deficits in pineal melatonin could suggest any protective effects occur earlier in stages of neoplastic development. For example, the increased expression of *Cdh2, Sema3f* and *Nrp1* would reduce shedding of cells from a tumor. This indicates a need to more extensively study *in vitro* properties (colony formation, wound healing, and trans-well cell invasion) and the time course of orthotopic allograft tumor development and metastasis, as well as incidence of TNBC induced by chemical carcinogens in our model.

### Clinical relevance

Exposure to LAN will reduce the duration and/or amount of melatonin action. Currently our data finds no *in vivo* effect of melatonin deficits, but *in vitro* gene expression does suggest melatonin could affect TNBC pathogenesis. If melatonin acts earlier in oncogenesis than invasion of circulating cells into tissue, such as reducing the prior shedding of cells from a tumor, then effective risk reduction would be best achieved with prophylactic reductions in ALAN exposure or use of melatonin supplements or agonists. Even if the risk reduction is modest, the prevalence and mortality of TNBC would make that effect meaningful.

## Contributions

CGD and PDS - *in vivo* experiments, analysis and paper preparation. DNT and SR - *in vitro* experiments, analysis and paper preparation. BAF, EOC and ZAS - in vivo experiments and analysis. JS and FDS - gDNA assay development and paper preparation. ELB - approach development and paper preparation. SDW - pinealectomy development. ST-conceived and led the study, conducted experiments and analysis, and led paper preparation.

